# Levels of p53 expression determine the competitive ability of embryonic stem cells during the onset of differentiation

**DOI:** 10.1101/2022.02.28.482311

**Authors:** Salvador Perez Montero, Sarah Bowling, Rubén Pérez-Carrasco, Tristan A. Rodriguez

**Affiliations:** National Heart and Lung Institute, Imperial College London, UK; Stem Cell Program, Boston Children’s Hospital, Boston, MA, USA and Department of Stem Cell and Regenerative Biology, Harvard University, Cambridge, MA, USA; Department of Life Sciences, Imperial College London, UK

**Author notes:** Both these authors contributed equally (Salvador Perez Montero, Rubén Pérez-Carrasco).

**Keywords:** cell competition, p53, mathematical modelling

## Abstract

During development, the rate of tissue growth is determined by the relative balance of cell division and cell death. Cell competition is a fitness quality control mechanism that contributes to this balance by eliminating viable cells that are less-fit than their neighbours. What mutations confer cells with a competitive advantage or the dynamics of the interactions between winner and loser cells are not well understood. Here, we show that embryonic cells lacking the tumour suppressor *p53* are super-competitors that eliminate their wild-type neighbours through the direct induction of apoptosis. This elimination is context dependant, as does not occur when cells are pluripotent and is triggered by the onset of differentiation. Furthermore, by combining mathematical modelling and cell-based assays we show that the elimination of wild-type cells is not through a competition for space or nutrients, but instead is mediated by short range interactions that are dependent on the local cell neighbourhood. This highlights the importance of the local cell neighbourhood and the competitive interactions within this neighbourhood for the regulation of proliferation during early embryonic development.

## INTRODUCTION

The regulation of growth and tissue homeostasis during embryogenesis relies on the proper balance of cell division and cell death. One mechanism contributing to this balance is cell competition. Cell competition is a quality control mechanism conserved from *Drosophila* to mammals that eliminates cells that although viable, are less fit than their neighbours. During competition those cells that are eliminated are generically termed losers. Accompanying this elimination, the fitter cells (winners) undergo compensatory proliferation, maintaining tissue homeostasis -reviewed in (Baker, 2020; Bowling et al., 2019; Diaz-Diaz and Torres, 2019; Johnston, 2014; Madan et al., 2018; Maruyama and Fujita, 2017; Vishwakarma and Piddini, 2020). For the purpose of this study, we define cell fitness as a cell’s ability to thrive in its environment. This ability is likely to be determined by a number of parameters, including signalling ability, metabolic rates and cell adhesion properties, but ultimately is established by the balance between the cell division rate and the sensitivity to cell death.

An important implication of cell competition is that cellular fitness is not only a cell-intrinsic property but is also determined relative to the fitness of neighbouring cells– a cell that is of sub-optimal fitness in one context may be ‘super-fit’ in the context of a different cell population. This is most clearly demonstrated in the case of wild-type cells, that in *Drosophila* and in mouse can eliminate a range of defective cells -reviewed in (Bowling et al., 2019), but can in turn also be eliminated by cells that over-express *Myc* (Claveria et al., 2013; de la Cova et al., 2004; Moreno and Basler, 2004; Sancho et al., 2013; Villa Del Campo et al., 2014). Due to this ability to induce the elimination of wild-type cells, cells over-expressing *Myc* have been termed super-competitors. The observation that cell fitness is relative to the cells neighbours implies that cells can interpret their relative fitness levels during cell competition. However, little is known regarding the mechanisms by which this process takes place. In *Drosophila* and cancer cells differential expression of *Flower* isoforms can act as fitness fingerprints for the cells winner and loser status (Madan et al., 2019; Rhiner et al., 2010). Similarly, in *Drosophila* innate immune-like signalling has been shown to induce for the elimination of *Minute* cells (that have a ribosomal deficiency) when they are surrounded by wild-type cells and for the elimination of wild-type cells by *Myc* super-competitors (Meyer et al., 2014). However, this pathway is not required for competition when the flies are maintained in a sterile environment (Germani et al., 2018) and promotes the overgrowth of polarity deficient cells when they are surrounded by wild-type cells (Katsukawa et al., 2018).

In mouse, at the onset of embryonic differentiation cell competition has been demonstrated to mediate the elimination of defective cells (Sancho et al., 2013) as well as those cells with low levels of *Myc* expression (Claveria et al., 2013). In the post-implantation embryo, just prior to gastrulation, this process eliminates around 35% of embryonic cells due to repression of the mTOR pathway (Bowling et al., 2018). This large-scale elimination is thought to ensure that only the fittest cells go on to contribute to further development and the germline (Bowling et al., 2019), and one important trigger of this elimination is mitochondrial dysfunction and mitochondrial DNA mutations (Lima et al., 2021). In the pre-implantation mouse embryo signalling via the Hippo pathway is required for the elimination through cell competition of mis-patterned cells (Hashimoto and Sasaki, 2019) and this pathway also mediates the elimination of normal human embryonic stem cells (ESCs) by karyotypically abnormal ones (Price et al., 2021), explaining how the abnormal cells expand in human pluripotent stem cell cultures.

Importantly, the cells eliminated in the early post-implantation mouse embryo display elevated P53 levels (Bowling et al., 2018; Lima et al., 2021) and cells with increased P53 expression are also eliminated during mouse during organogenesis (Zhang et al., 2017). In Madin-Darby Canine Kidney (MDCK) cells, mutation of the cell polarity gene *Scribble* in a mosaic fashion leads to an increase in P53 that induces mutant cell elimination via mechanical cell competition (Wagstaff et al., 2016). The observation that in mouse chimeras, *p53* mutant cells have a selective advantage during development (Dejosez et al., 2013), suggests that loss of P53 expression provides cells with a competitive advantage. A further fascinating indication of the role of P53 in competition comes from analysing interspecies chimeras as well as co-cultured human and mouse pluripotent stem cells. Both in vivo and in vitro, human ESCs are eliminated by mouse ESCs by cell competition. However, mutation of *p53* in the human cells is sufficient to prevent the human ESC elimination (Zheng et al., 2021). Together, these data suggest that relative P53 levels are a key determinant of embryonic fitness. Here we have asked if in mouse, mutation of *p53* turns cells into super-competitors. By comparing the behaviour of wild-type and *p53* mutant cells in separate versus coculture conditions and by developing a mathematical model of this interaction, we demonstrate that *p53* null mutant mouse ESCs actively induce the apoptotic elimination of wild-type cells in co-culture. Furthermore, we also find that this elimination is mediated by short-range signalling, highlighting the importance of local competitive interactions for the regulation of cell proliferation during the onset of differentiation.

## RESULTS

### P53 mutant ESCs behave as super-competitors

The observation that in the embryo and in ESC models of cell competition loser cells show an increase in P53 expression (Bowling et al., 2018; Lima et al., 2021) combined with the finding that in chimeras *p53* mutant cells contribute preferentially to the embryo (Dejosez et al., 2013) suggests that differences in the levels of P53 expression determine the competitive ability of embryonic cells. To test this possibility, we generated *p53* null mutant mouse ESCs (Fig. 1A) and compared their behaviour when they were cultured in a homogeneous (separate) culture to when they were co-cultured with wild-type cells. We found that in pluripotency culture conditions, *p53* mutant cells grew at a similar rate to cells both in separate and co-culture conditions (Fig. 1B-C). Here 8×10^5^ cells of each genotype were plated in separate culture and the co-culture contained a mix of 4×10^5^ cells of each genotype, so also totalling 8×10^5^ cells. However, in contrast to this, when the same cell numbers were plated and the ESCs were induced to differentiate by culture in N2B27, *p53*^*-/-*^ cells displayed a small proliferative advantage in separate culture and induced the growth arrest of wild-type cells in co-culture (Fig. D-E). This behaviour was not due to the inability of *p53* mutant cells to initiate differentiation, as when cultured in N2B27 *p53*^*-/-*^ cells down-regulate the expression of naïve pluripotency markers and increase the expression of the post-implantation epiblast marker *Fgf5* in a similar way as control cells do (Fig. 1F). These results suggest that mutation of *p53* makes cells into super-competitors that out-compete wild-type cells.

**Figure 1.**
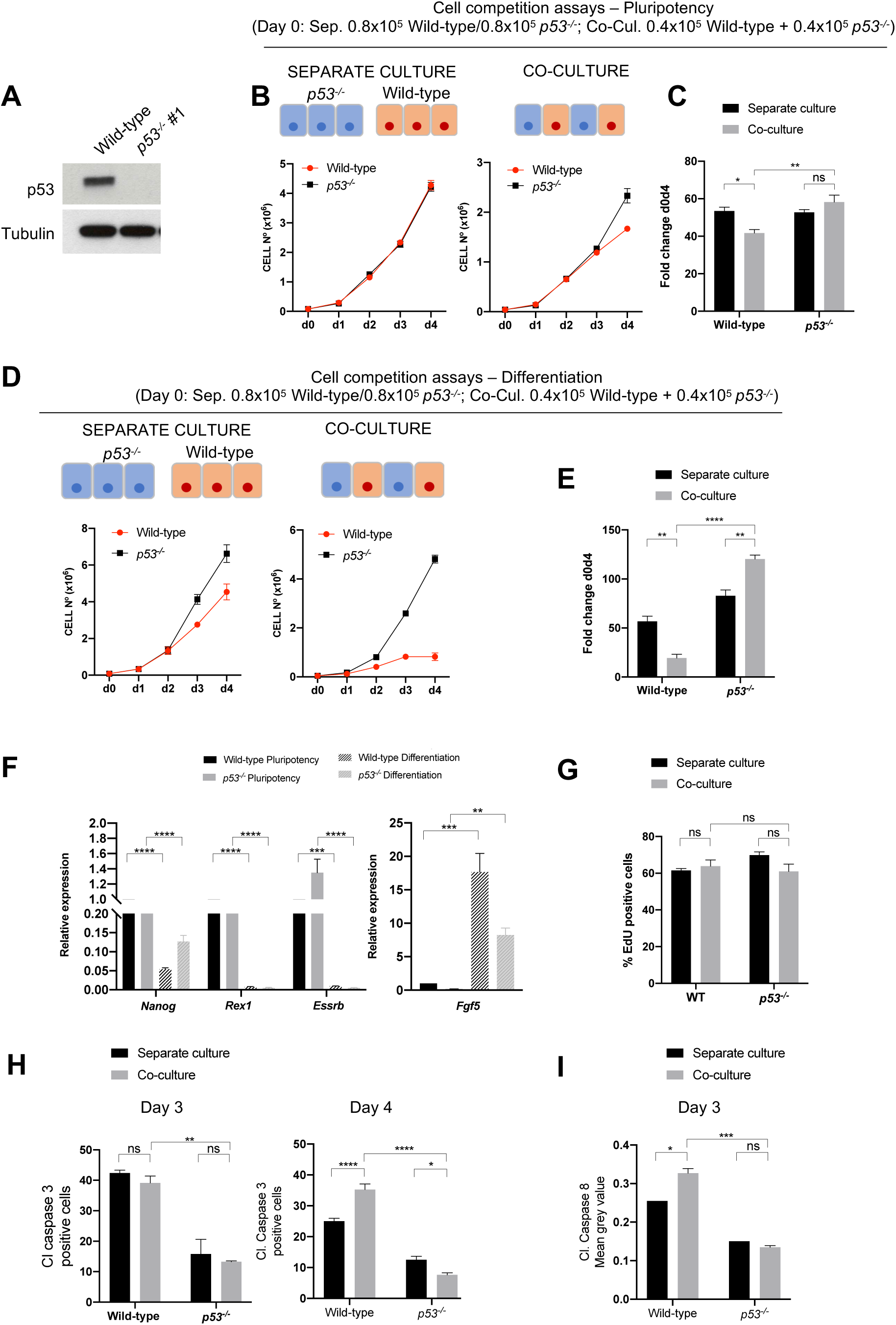
Wild-type cells are outcompeted by *p53*^*-/-*^ cells. **A**. p53 levels in wild-type and two different *p53*^*-/-*^ clones untreated or treated with the p53 activator Nutlin-3a for 4h. **B**. Growth curves of wild-type and *p53*^*-/-*^ cells over 4 days in separate or co-culture in pluripotency conditions. **C**. Fold change in wild-type and *p53*^*-/-*^ cell numbers between day 0 and day 4 when cultured alone or co-cultured in pluripotency conditions. **D**. Growth curves of wild-type and *p53*^*-/-*^ cells over 4 days in separate or co-culture differentiation conditions. **E**. Fold change in wild-type and *p53*^*-/-*^ cell numbers between day 0 and day 4 when cultured alone or co-cultured in differentiation conditions. **F**. Quantitative RT-PCR showing gene expression levels of naïve and primed pluripotency markers in wild-type and *p53*^*-/-*^ ESCs in pluripotency and differentiation. Gene expression is normalized against beta-Actin. **G**. Percentage of EDU incorporation in wild-type cells and *p53*^*-/-*^ cells cultured for 4 days separately or co-cultured. **H**. Percentage of cleaved caspase 3 positive cells in wild-type and *p53*^*-/-*^ cells cultured for 3 and 4 days separately or co-cultured. **I**. Levels of Cleaved Caspase 8 in Wild-type and *p53*^*-/-*^ cells cultured alone or co-culture for 3 days. Data were obtained from three independent experiments and are shown as the mean + SEM (**B**-**I**).

The cyclin dependant kinase inhibitor *Cdkn1a* (*p21)* is an important target activated by P53 (Pilley et al., 2021). To test if P21 regulation of the cell cycle is an important part of the mechanism by which P53 confers a super-competitor status we generated *p21* mutant ESCs (Fig. S1A). We found that when these cells were cultured in N2B27 they could differentiate normally (Fig. S1B) and grew similarly to control cells in both separate and co-culture conditions (Fig. S1C-D), and therefore do not show super-competitor behaviour. The observation that both *p53*^*-/-*^ and *p21*^-/-^ cells displayed similar levels of Edu incorporation to control cells in both separate and co-culture with control cells (Fig. 1G and Fig. S1E) further supports the conclusion that regulation of the cell cycle is not the primary mechanism by which *p53* mutant cells become super-competitors.

Given that we did not find evidence for differences in the cycle explaining the increased competitive ability of *p53* mutant cells we investigated the role of the apoptotic response. When we analysed cleaved-caspase 3 expression we observed that it was significantly lower in *p53*^*-/-*^ ESCs when compared to control cells after 3 days and after 4 days culture in N2B27 (Fig. 1H). This indicates that *p53* mutant cells are intrinsically more resistant to apoptosis than wild-type cells and likely explains their faster growth rate in separate culture. Interestingly, we also observed that wild-type cells at day 4 of co-culture with *p53*^*-/-*^ ESCs showed an increase in cleaved-caspase 3 expression compared to when they were maintained in a homotypic (separate) culture (Fig. 1H). The fact that this difference was not apparent at day 3 of co-culture, even though at this time-point there is a clear difference in the growth rate of *p53*^*-/-*^ and control ESCs, suggests that there may be caspase 3 independent death occurring before this stage. To address this possibility, we analysed the expression of cleaved-caspase 8, that mediates extrinsic cell death signalling (Fuchs and Steller, 2015). We not only found that *p53*^*-/-*^ cells displayed lower levels of cleaved caspase-8 expression than wild-type cells in separate culture (Fig. S2A-B), but also that at day 3 wild-type cells showed higher cleaved-caspase 8 expression in co-culture compared to separate culture (Fig. 1I and Fig. S2C). These results suggest that *p53*^*-/-*^ cells out-compete wild-type cells by inducing their apoptotic elimination.

### Developing a mathematical model of super-competition

Our results indicate that *p53* mutant cells outcompete wild-type cells in coculture. To gain further insight into this competition and explore the mechanism by which loser cell elimination takes place, we developed a mathematical model to recapitulate quantitatively the differential cell population dynamics in separate and co-culture assays. The aim of this model is twofold: on one hand disentangle potential confusing effects of population intrinsic growth and cell-cell competition effects on total cell population growth, while at the same time obtain mechanistic information of the nature of competition. We described the evolution in time of the number of wild-type cells (*W*) and *p53*^*-/-*^ cells (*P*) by specifying a set of ordinary differential equations (ODE) describing the effect that different population compositions (W, P) have on the net growth of each species. In the absence of competition – at low cell numbers – each species population grows with an intrinsic rate *ρ*_*i*_ > 0, *i* = {*W, P*} (Fig. 2A top), this is the net growth of the population incorporating the intrinsic proliferation and apoptotic rates. As the population grows, cells compete with each other with a strength *k*_*j*_ specific for each of the 4 possible interactions (*j* = {*WW, WP, PW, PP*}, see Fig 3A-B),

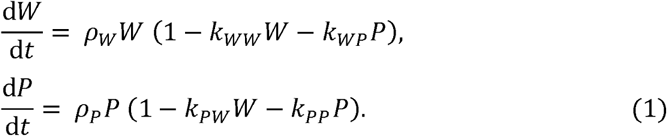

This Lotka-Volterra description has been successfully used in a different context of cell competition (Ram et al., 2019) and assumes that the magnitude of the competition is proportional to the number of cells of each type in the dish (or equivalently, their concentration). Therefore, the increase in apoptotic rate in a crowded dish is translated into a decrease of the total net growth rate of the population of each species, making it negative in crowded environments (e.g. 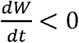, see Fig. 2A). To test the model, we used Bayesian inference to identify which portion of the parameter space {*p*_*i*_, *k*_*j*_} is consistent with the experimental data by using a likelihood function that compares experimental and numerical trajectories (see Materials and Methods)(Toni et al., 2009). The result of this analysis is a preliminary credibility distribution of the parameters of the model compatible with the cell population trajectories shown in Fig. 1. Inspection of these distribution of parameters allowed us to identify which additional experimental initial plating conditions would carry information to improve our predicted parameter distributions. Experiments were then performed using these identified plating conditions and were analysed in the same manner as described above. This resulted in an iterative analysis composed of a set of experimental trajectories for 24 different plating conditions that generated a posterior credibility distribution of the parameters that can be used to extract mechanistic information of the competition (Fig. S3). The trajectories corresponding to this distribution show a very good fit to the model (Fig. 2C and Fig. S4). Analysis of the pairwise relationship between parameters confirm that the intrinsic net growth of *p53*^*-/-*^ is faster than the intrinsic net growth of wild-type cells, *ρ*_*P*_ > *ρ*_*W*_ (Fig. 2D). In addition, comparison of the competition strengths revealed that there were not significant differences between the competition of *p53*^*-/-*^ cells with themselves compared to the corresponding homotypic competition between wild-type cells (*k*_*WW*_ ≃ *k*_*pp*_). On the other hand, as expected, the presence of *p53*^*-/-*^ ESCs induced a dramatic decrease on the growth rate of wild-type cells (*k*_*WW*_ < *k*_*WP*_).Strikingly, *p53* ESC growth was unaffected by the identity of the cells in their neighbourhood (*k*_*PP*_ ≃ *k*_*PW*_). Our model therefore quantified the interactions between cells allowing us to identify differential cellular properties and compare their magnitude.

**Figure 2.**
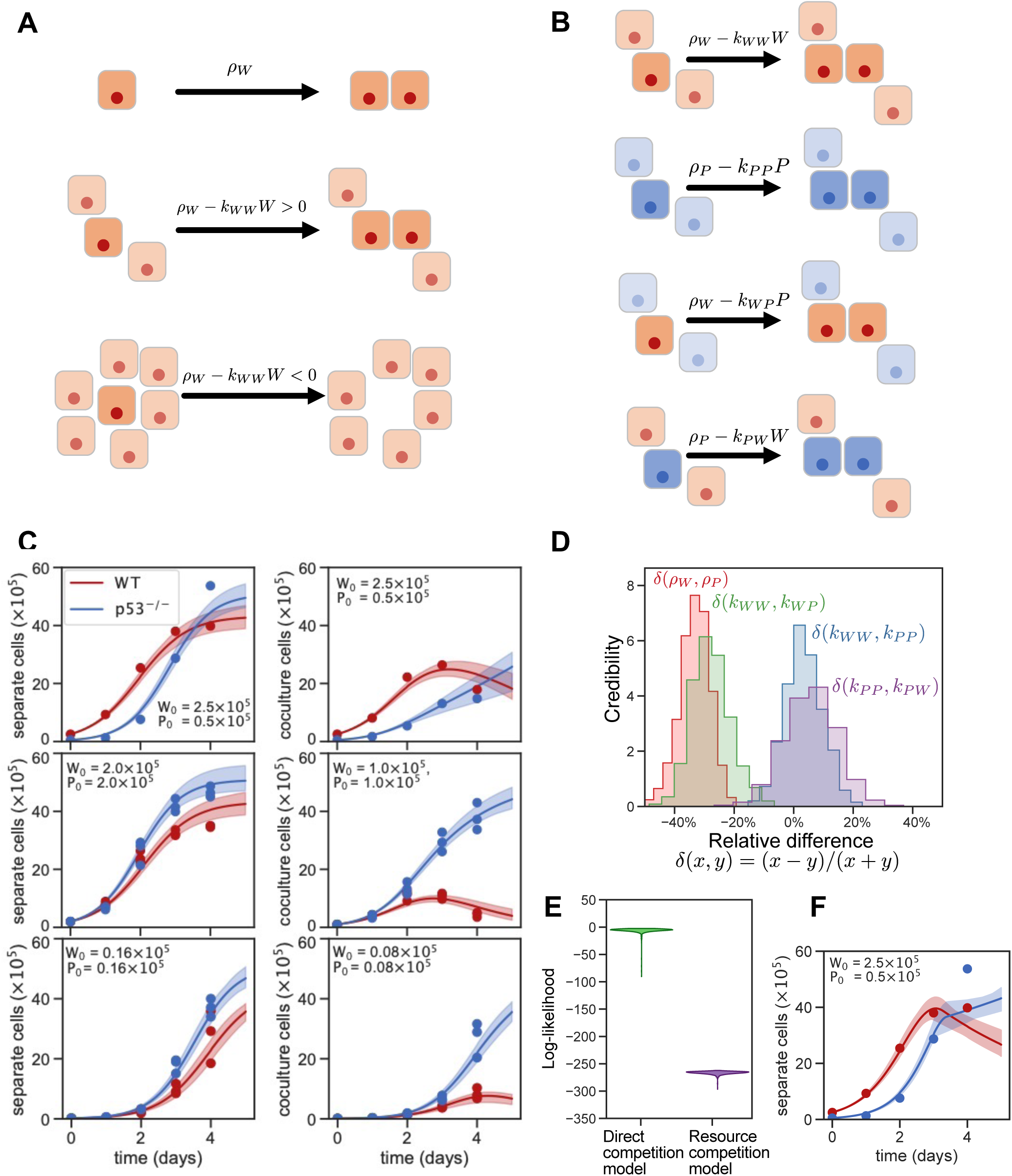
A mathematical model suggests a direct asymmetric competition. **A** Schematic depicting how the net proliferation rate per cell in a homotypic population depends on cell density. **B** Schematic showing the four possible direct competition mechanisms affecting population growth of WT and *p53*^*-/-*^ cells. **C** Comparison of the experimental data (one circle per replicate) with the model prediction (lines) for the direct competition model (Eq. 1) for 9 of 24 the initial conditions used in the inference (the rest can be found in Fig. S3). Shaded zones show model prediction for the inferred parameter region with likelihood >90% of its maximum. Parameter inference values can be found in Fig. S4. **D** Inferred credibility distributions for the relative differences between different model parameters. Relative differences are significant between the intrinsic growths and between the competition constants on WT cells. **E** Log-likelihood distributions of the direct competition model (Eq. 1) and the resource competition model (Eq. 3). **F** Comparison of the experimental data with the resource competition model, visualization details are the same as in panel C. Parameter inference values can be found in Fig. S4.

**Figure 3.**
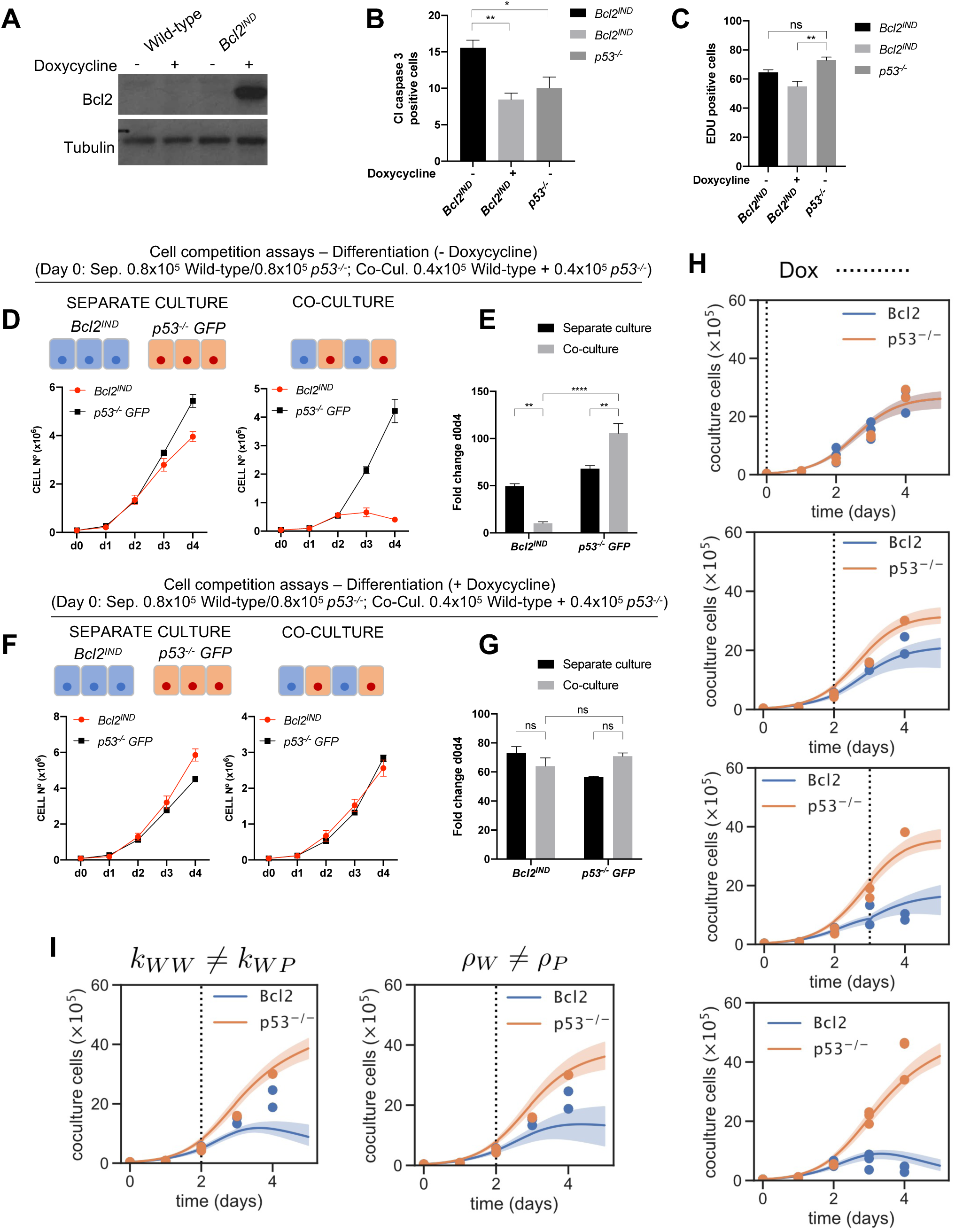
*p53*^*-/-*^ ESCs induce the apoptotic elimination of wild-type cells in coculture. **A**. Bcl2 levels in wild-type and *Bcl2*^*IND*^ cells not treated or treated with doxycycline for 3 days. **B**. Percentage of cleaved caspase 3 positive cells in *Bcl2*^*IND*^ treated with or without doxycycline and *p53*^*-/-*^ cells. **C**. Percentage of EDU incorporation in *Bcl2*^*IND*^ treated with or without doxycycline and *p53*^*-/-*^ cells. **D**. Growth curves of *Bcl2*^*IND*^ and *p53*^*-/-*^ cells over 4 days in separate or co-culture conditions without doxycycline treatment. **E**. Fold change in wild-type and *p53*^*-/-*^ cell numbers between day 0 and day 4. **F**. Growth curves of *Bcl2*^*IND*^ and *p53*^*-/-*^ cells over 4 days in separate or co-culture conditions with doxycycline treatment from day 1. **G**. Fold change in wild-type and *p53*^*-/-*^ cell numbers between day 0 and day 4. Data were obtained from three independent experiments and are shown as the mean + SEM (**E**-**H**). **H** Comparison of the direct competition model (lines) with the experimental data (one replicate per circle) for doxycycline treatment at different days (vertical dashed lines). Shaded zones show model prediction for the inferred parameter region with likelihood >90% of its maximum. Parameters are the same as in Figs. 2, S3, S4. **I** Resulting model trajectories when doxycycline treated cells are simulated by only changing the intrinsic growth and maintaining differences in competition strength (left), or only changing the competition strength and maintaining differences in intrinsic growth (right).

### *P53* mutant cells induce the apoptotic elimination of wild-type ESCs

Our model indicates that *p53*^*-/-*^ ESCs have a direct negative effect on wild-type cells and the increased expression of apoptotic markers in cocultured wild-type cells (Fig. 1H-I and S2A-C) suggests that this takes place through the induction of their apoptotic elimination. To test this possibility, we first performed the cell competition assays in the presence of a pan-caspase inhibitor (Z-VAD-FMK). For this we cultured wild-type and *p53*^*-/-*^ ESCs in N2B27 in separate and co-culture conditions and added the caspase inhibitor from day 2 to day 4 of culture. We found that this partially rescued the elimination of wild-type cells in co-culture (Fig. S2D), raising the prospect that non-apoptotic forms of cell death may be contributing to the out-competition of wild-type cells.

BCL-2 is a key anti-apoptotic protein acting in the mitochondrial apoptotic pathway (Tait and Green, 2013) and *Bcl-2* over-expression prevents loser cell elimination in interspecies chimeras (Zheng et al., 2021). We therefore generated doxycycline inducible ESCs (Fig. 3A). Induction of BCL2 expression by doxycycline addition decreased the intrinsic apoptotic rate to levels that were similar to those found in *p53* mutant cells (Fig. 3B). In contrast to this BCL2 induction did not significantly affect proliferation rates (Fig. 3C). We therefore assayed the behaviour of *Bcl-2* inducible ESCs (*Bcl2*^*Ind*^) in separate and co-culture with *p53*^*-/-*^ cells. We found that similarly to what occurred with wild-type ESCs, when *Bcl2*^*Ind*^ ESCs were cultured separately in N2B27 without the addition of doxycycline they grew slightly slower than *p53*^*-/-*^ ESCs and when co-cultured with *p53*^*-/-*^ ESCs they were effectively eliminated (Fig. 3D-E). In contrast to this, we observed that upon BCL2 induction with doxycycline *Bcl2*^*Ind*^ ESCs grew similarly to *p53*^*-/-*^ cells, both in separate and co-culture condition (Fig. 3F-G) and are therefore no longer eliminated by *p53*^*-/-*^ cells. These results indicate that *p53* mutant cells out-compete wild-type cells by inducing their apoptotic elimination.

To analyse the dynamics by which BCL2 rescues the competition, we used our model together with the inferred parameters to reproduce experimental trajectories of doxycycline induced rescue at different timepoints of the competition. The model successfully reproduced the dynamics of the rescue (Fig. 3H). Untreated Bcl2^*Ind*^ ESCs were successfully simulated using the same parameters as wild-type cells. In contrast to this, to reproduce the experimental trajectories of Bcl2^*Ind*^ cells treated with doxycycline required to eliminate completely the asymmetry in competition, reducing the competition strength that *p53* mutant cells have on Bcl2^*Ind*^ (*k*_*WP*_) to the same level than homotypic competition (*k*_*WW =*_ *k*_*WP*_) (Fig. 3I). Interestingly, it also required to eliminate the asymmetry in the intrinsic population growth by increasing the net proliferation rate of *Bcl2*^*Ind*^ cells to the same level as *p53*^*-/-*^ ESCs (*ρ*_*W =*_ *ρ*_*P*_). In summary, BCL2 rescue dynamics could only be reproduced by the model when all the parameters of wild-type cells are restored to the equivalent parameter of *p53*^*-/-*^ ESC, supporting the hypothesis that the competition mechanism is an active elimination of wild-type cells by *p53*^*-/-*^ cells.

### Short range signalling mediates the elimination of wild-type cells by *p53*^-/-^ ESCs

The data presented in Figs. 1-3 points to a direct competition between *p53*^*-/-*^ and wild-type cells, whereby one induces the elimination of the other. But in addition to the direct competition model (Eq. (1)) we also wanted to explore the possibility that cells compete for a shared resource that is consumed over time with differential rates and tolerances depending on the cell type (see Materials and Methods, Eq. (3)). Interestingly, this model was not able to recapitulate the experimental growth curves (Fig. 2E-F and Fig. S3). To test this assumption experimentally we analysed the effect that providing unlimited amounts of nutrients has on the dynamics of the competition between wild-type and *p53*^*-/-*^ cells. For this we performed our competition experiments in a culture media that was continuously perfused from day 1 of culture by a pump (Fig. 4A). Using this system, we observed that both wild-type and *p53*^*-/-*^ ESCs grew roughly double as much when the media was perfused compared to when it wasn’t, in both separate and co-culture conditions (Fig. 4B). This increase in growth was likely due to a lower level of cell death, as the percentage of cells that were positive for cleaved caspase 3 was also significantly reduced in the perfusion condition (Fig. 4C). It is also possible that the apoptotic cells are being more efficiently removed due to the perfusion. Notably, we found that despite these lower levels of apoptosis, the perfusion did not significantly affect the degree to which wild-type ESC were eliminated in co-culture (Fig. 4D)., suggesting that the latter may be true. These results suggest two things. First, they indicate that the elimination of wild-type cells by *p53*^*-/-*^ ESCs is unlikely to be due to nutrient depravation. Second, the fact that the number of cells in the culture dish can double without increasing the degree of wild-type apoptosis, also suggests that this elimination is not due to mechanical stress, as has been shown to be the case for MDCK cells with increased p53 expression (Wagstaff et al., 2016).

**Figure 4.**
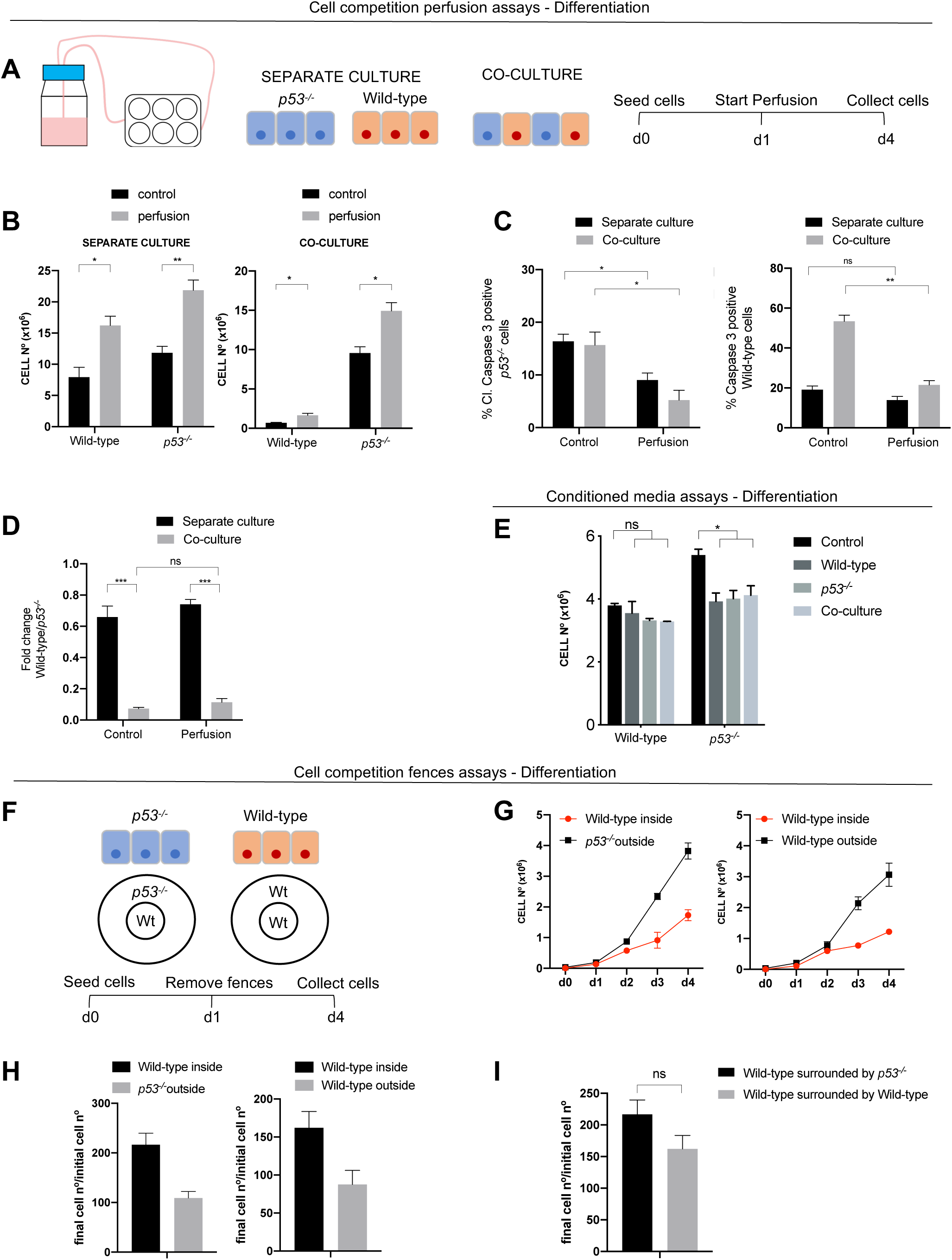
Wild-type cell elimination is cell contact dependent. **A**. Schematic experimental setup whereby cells were cultured separately and together and N2B27 media was perfused over the plate from day 2 until day 4. Cells were then counted and fixed. **B**. Wild-type and *p53*^*-/-*^ cell numbers over 4 days cultured separately or together in a control plate and in a perfused plate. **C**. Fold change in wild-type cell numbers between day 0 and day 4 when cultured alone or with *p53*^*-/-*^ cells in N2B27 in a control plate and in a perfused plate. **D**. Percentage of cleaved caspase 3 in wild-type and *p53*^*-/-*^ cells cultured separately or in co-culture with and without perfusing N2B27 media. **E**. Cell numbers of wild-type and *p53*^*-/-*^ cells cultured separately for 4 days and adding conditioned media from day 3 to day 4. Conditioned media was taken from cultured wild-type and *p53*^*-/-*^ cells or from a co-culture of both cells types. **F**. Schematic experimental setup whereby wild-type cells where cultured surrounded by *p53*^*-/-*^ cells or wild-type cells in the same well without cell contact between both cell types. **G**. Growth curves of wild-type and *p53*^*-/-*^ cells over 4 days cultured as in F. **H**. Fold change in wild-type and *p53*^*-/-*^ cell numbers between day 0 and day 4. **I**. Fold change in wild-type cell numbers a between day 0 and day 4 when they are cultured surrounded by wild-type or *p53*^*-/-*^ cells. Data were obtained from three independent experiments and are shown as the mean + SEM (**A**-**I**).

We next analysed the possibility that diffusible growth factors secreted into the media could be inducing the elimination of loser cells. For this we did two things. First, we analysed the effect of conditioned media taken from wild-type and *p53* mutant cells cultured separately, as well as well as taken from their co-culture. We observed that none of these conditioned medias had any effect on the growth of wild-type cells and they all reduced *p53*^*-/-*^ cell growth (Fig. 4E). To further address the importance of secreted factors we used a fences system (Lawlor et al., 2020), whereby one population of cells is grown surrounded by another but separated by fences that are removed once the cells are seeded (Fig. 4F). This allows the different cell populations to be cultured without contact but sharing the same media. When this was done, we observed that cells grown on the outside layer grew slower than those cultured on the inside, irrespective of their genotype (Fig. 4G-H). Importantly, we found that wild-type cells grew similarly if they were surrounded without contact by *p53* mutant cells or by other wild-type cells (Fig. 4I). Together this data suggest that cell-cell contact or short-range signalling is required for *p53*^*-/-*^ cells to eliminate wild-type cells.

### Cell neighbourhood changes during cell competition

Given the importance of short-range signalling for the elimination of wild-type cells, we asked if changes in cell neighbourhood could explain the competition between wild-type and *p53* mutant cells. We consider here the local cell neighbourhood to be the relative numbers of wild-type and *p53*^*-/-*^ cells that are in direct contact with a given cell in the co-culture condition. When we analysed the cell neighbourhood of wild-type cells, we found that during the time-course of the experiment these cells increased the average number of wild-type neighbours until reaching a peak at day 2 of co-culture, but that after this time-point this number of wild-type neighbours decreased (Fig. 5A-B). In contrast to this, the average number of *p53*^*-/-*^ neighbours that wild-type cells had increased from day 1 and by day 3 they had more *p53*^*-/-*^ neighbours than wild-type neighbours. This switch in wild-type neighbourhood between days 2 and 3 coincided with when wild-type elimination is most obvious in the growth curves of these cells (Fig. 1C). The same neighbourhood dynamics could be observed for *Bcl2*^*Ind*^ cells when these were co-cultured with *p53*^*-/-*^ ESCs without doxycycline (Fig. 5C). However, if BCL2 is induced with doxycycline and loser cell elimination is prevented, then *Bcl2*^*In*^ cells showed a daily increase in *Bcl2*^*In*^ cell neighbours, while the number of *p53*^*-/-*^ ESC neighbours first increased until day 2 and then decreased (Fig. 5D). These results are consistent with cell neighbourhood having a role in the outcome of competition.

**Figure 5.**
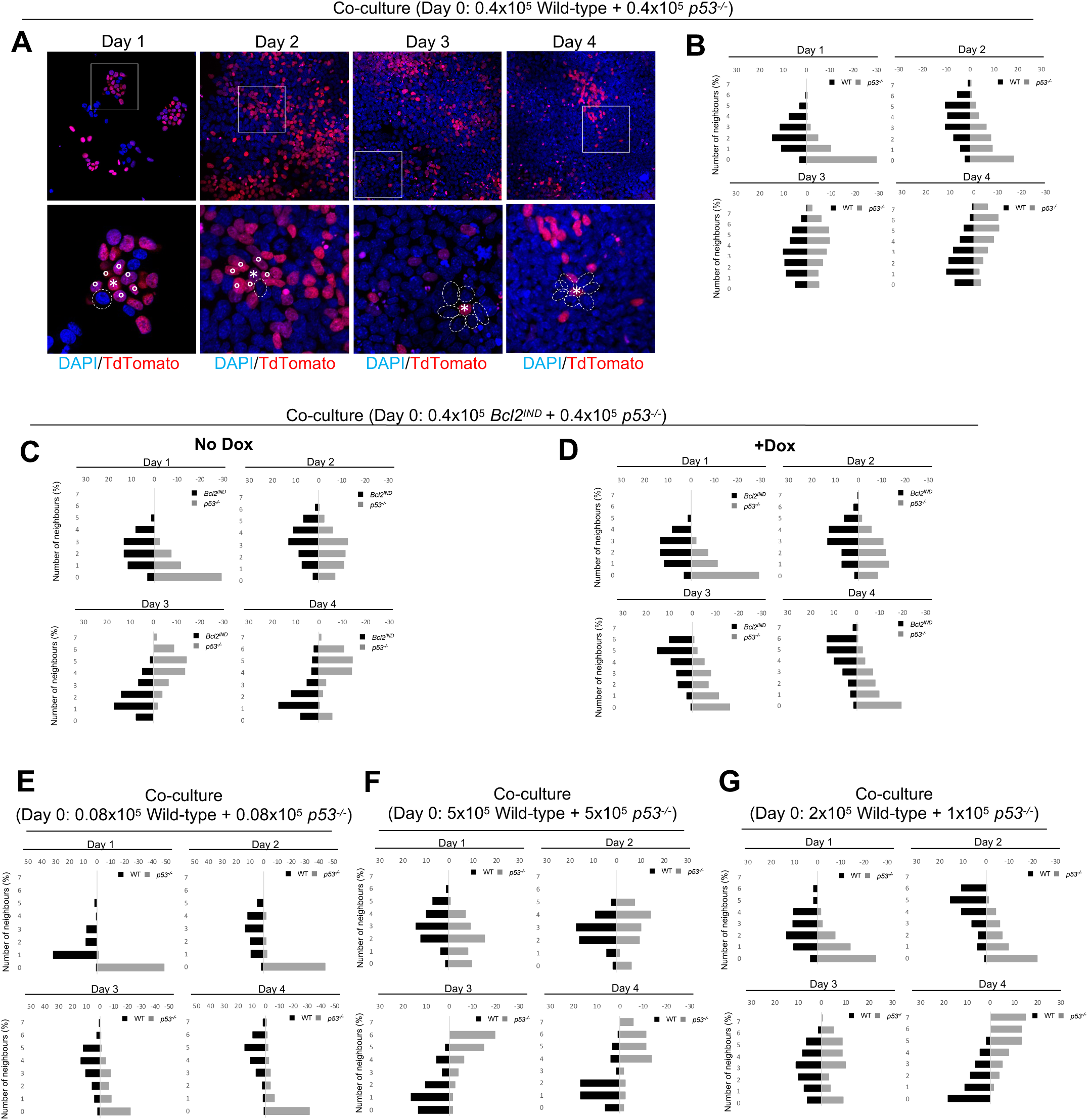
Neighbourhood changes during cell competition. **A**. Immunostainings of wild-type and *p53*^*-/-*^ cells co-cultured from day 1 to day 4 of the culture. An example of neighbours of wild-type cells is shown for each day. **B**. Number of wild-type and *p53*^*-/-*^ neighbours of wild-type cells from day 1 to day 4 of the co-cultures. **C**. Number of *Bcl2*^*IND*^ and *p53*^*-/-*^ neighbours of *Bcl2*^*IND*^ cells from day 1 to day 4 of co-cultures without doxycycline treatment. **D**. Number of *Bcl2*^*IND*^ and *p53*^*-/-*^ neighbours of *Bcl2*^*IND*^ cells from day 1 to day 4 of co-cultures with doxycycline treatment from day 1. **E**. Number of wild-type and *p53*^*-/-*^ neighbours of wild-type cells from day 1 to day 4 of co-cultures with lower initial cell numbers plated. **F**. Number of wild-type and *p53*^*-/-*^ neighbours of wild-type cells from day 1 to day 4 of co-cultures with higher initial cell numbers plated. **G**. Number of wild-type and *p53*^*-/-*^ neighbours of wild-type cells from day 1 to day 4 of co-cultures with different percentage of seeded cells. Data were obtained from three independent experiments and are shown as the mirrored distribution of wild-type and *p53*^*-/-*^ neighbour numbers (**B**-**G**).

We next tested the effects of changing the concentrations of cells plated. First, we analysed the effects of reducing the cell numbers plated. For this a mix of 0.08×10^5^ wild-type cells and 0.08×10^5^ *p53*^*-/-*^ ESCs were plated in the co-culture condition. We observed that at this low confluency there was no substantial change in the neighbourhood of wild-type cells in co-culture (Fig. 5E). Interestingly, this correlated with the total number of wild-type cells still increasing at day 4 of co-culture (Fig. S5A). This suggests that the low confluency is reducing the number of possible *p53*^*-/-*^ neighbours, and that this is decreasing the rate of competition. We then analysed the effects of increasing the cells numbers plated at day 0 to a mix of 5×10^5^ wild-type and 5×10^5^ *p53*^*-/-*^ cells in co-culture. This condition led to wild-type cells having more *p53*^*-/-*^ cell neighbours from day 2 (Fig. 5F) and being robustly eliminated in co-culture (Fig. S5B). Similar was observed when we changed the proportion of wild-type to *p53*^*-/-*^ cells plated. When a mix of 2×10^5^ wild-type and 1×10^5^ *p53*^*-/-*^ cells were plated in co-culture this caused a shift between days 2 and 3 in the neighbourhood of wild-type cells from having more wild-type neighbours to having more *p53*^*-/-*^ ones (Fig. 5G). This correlated with when the wild-type cells were eliminated in co-culture (Fig. S5C). These results, together with the importance of short-range signalling identified in Fig. 4, point to the direct interaction between wild-type and *p53*^*-/-*^ cells determining the outcome of their competition. Our findings also suggest that the relative number of neighbours with a different competitive ability is a factor affecting the replacement of wild-type cells.

## DISCUSSION

Cell competition is a fitness quality control mechanism that eliminates cells that are less-fit than their neighbours. One important implication of cell competition is that the fitness of a cell is relative to the fitness of its neighbour. For example, in mouse, wild-type cells eliminate cells with mitochondrial dysfunction (Lima et al., 2021), but are eliminated by cells that over-express c-MYC (Claveria et al., 2013; Sancho et al., 2013). Here, we have addressed how the expression levels of P53 affects the competitive nature of pluripotent cells. We find that cells lacking *p53* behave as super-competitors and eliminate their wild-type neighbours. This elimination is dependent on the onset of differentiation, as when cells are cultured in pluripotency conditions mutation of p53 provides cells with a competitive advantage. Furthermore, by combining mathematical modelling and cell-based assays we also show that the competitive advantage of *p53* null mutant cells is mediated by short-range signalling interactions that induce the elimination of wild-type cells, rather than through a competition for nutrients or space. This highlights the importance of the local cell neighbourhood for the regulation of proliferation during early embryonic development.

Over the last few years there has been increasing evidence for a key role for P53 in cell competition. In *Drosophila* p53 has been shown to be required in Myc-overexpressing super-competitor cells to induce wild-type cell elimination (de la Cova et al., 2014). In these winner cells, p53 regulates metabolism by promoting oxidative phosphorylation and inhibiting glycolysis. In contrast to this, mechanical stress induces increased *p53* expression in polarity deficient MDCK cells and this activation causes the elimination of these loser cells (Wagstaff et al., 2016). In mouse cell competition, p53 has been shown to play multiple roles. In haemopoietic stem/progenitor cells, DNA damage causes the out-competition of cells with higher p53 levels (Bondar and Medzhitov, 2010). Here the damage induces senescence in those cells that have higher p53 levels than their neighbours. In the mouse embryo it was found that clones carrying a mutation of *Mdm2/4*, and therefore that have elevated P53 expression, are outcompeted by wild-type cells (Zhang et al., 2017). Similarly, we have shown that during the onset of differentiation, defective cells such as cells with impaired BMP signalling or tetraploid ESCs, show increased p53 expression that is required for their out-competition by wild-type cells. We also demonstrated that the mechanism of elimination of these defective cells is because in a competitive environment p53 represses mTOR signalling and this induces apoptosis. Our results, together with these studies, suggests therefore that p53 could act as a general sensor of cell fitness across different tissues.

Three main modes of cell competition have been proposed, competition for nutrients, mechanical competition, and direct fitness sensing between cells (Bowling et al., 2019). For example, during cell competition in MDCK cells, elevated p53 expression sensitises cells to compaction, indicating that differences in p53 expression determine a differential response to mechanical stress (Wagstaff et al., 2016). In *Drosophila*, p53 regulates cell metabolism to determine the competitive nature of cells in the imaginal wing disc (de la Cova et al., 2014). This metabolic role could be used to infer that p53 is required to establish differences in nutrient metabolism that will in turn direct the outcome of competition. But our studies described here analysing the behaviour of *p53* mutant pluripotent cells suggest that during ESC differentiation p53 is possibly playing roles that are different to those described above for MDCK cells and in *Drosophila*. Our observation that there is no change to the cell competition dynamics between wild-type and *p53*^*-/-*^ ESCs despite cell numbers doubling when assays are performed in media that is being continuously replenished through cell perfusion, suggests that neither nutrient or space availability is determining the outcome of the competition between these cell types. Instead, our findings reveal that a change in the cell neighbourhood of loser wild-type cells is correlated with the outcome of cell competition. This suggests that the local interaction between wild-type and mutant cells regulates the elimination of the wild-type cells. These results are in line with what has been observed with *c-Myc* over-expressing ESCs, that eliminate wild-type cells via short-range/contact dependant signalling (Diaz-Diaz et al., 2017) and contrast with our own findings that the elimination of ESCs with defective BMP signalling is mediated by long-range signalling (Sancho et al., 2013). Our observations that the relative number of winner/loser neighbours determines the outcome of cells competition are also in accordance with what has been found in the *Drosophila* pupal notum, where the super-competition ability of *c-Myc* over-expressing cells has been related to the relative surface area shared between winner and loser cells (Levayer et al., 2015). This suggests that replacement of wild-type cells in a tissue occurs through a different mechanism than the replacement of dysfunctional cells, and therefore during embryonic development there may be several distinct forms of cell competition acting in a tissue at the same time.

Computational and mathematical modelling has provided invaluable insight allowing us to distinguish between possible models of competition, as well as revealing and quantifying the minimal rules required to reproduce the competition dynamics observed. Further descriptions of local competition in confluent populations – where the population cannot be considered homogeneous - will require a mathematical framework that incorporates the spatial dynamics of the competition. Such models can use similar ODE formulations where clone shape is taken into account implicitly in the functional form of the competition terms (Nishikawa et al., 2016). Alternatively, more detailed descriptions of the cellular monolayer can be simulated by using a computational vertex model, where the dynamics and environment of each individual cell is taken into account. These spatial models can allow us to study additional modes of competition resulting from dynamics exclusive to confluent tissues. For example, interaction between cell populations growing at different rates has been shown to introduce a mechanical stress that can act as a feedback mechanism to stabilize uniform growth (Shraiman, 2005). Similarly, cell geometry and mechanical heterogeneities within a developing tissue can bias mechanical cell elimination leading to effective changes in competition properties of coexisting cellular populations (Lee and Morishita, 2017). In addition, following cell elimination, cell-type specific topological remodelling of epithelial junctions has been shown to be enough to induce a difference in the fitness between cell populations in the *Drosophila* wing disc (Tsuboi et al., 2018).

More sophisticated agent-based multi-scale models can be used to simulate more specific cell and tissue morphologies, such as competition during the transition to congruence. An example of such models was used in (Gradeci et al., 2021) using automatic annotation of movies of co-cultured wild-type and polarity deficient MDCK cells lasting up to 4 days, providing extensive input on the behaviour of wild-type and mutant cells. Their modelling identified that cell density and stiffness is sufficient to account for the apoptotic elimination of loser cells during mechanical competition. In contrast to this, the outcome of biochemical competition appears to be primarily regulated by the organization of winner and loser cells in the tissue. Interestingly, these results are very much in accordance with the likely role of cell neighbourhood identified in our study. One of the main limitations of the successful application of spatial computational models is the large amount of parameters and rules that can be used to describe the behaviour of cellular populations, making hard to infer confidently details of the model even in simple scenarios (Kursawe et al., 2018). This highlights the necessity of minimal models able to incorporate and test mechanistic hypothesis matching the complexity of the available data.

In conclusion, our studies identify that upon exit from pluripotency, loss of p53 expression is sufficient to make ESCs into super-competitors. These winner cells induce the replacement of healthy wild-type cells via a direct induction of apoptosis. This cell replacement not only provides a potential explanation for the expansion of cells with p53 mutations in human pluripotent stem cell cultures (Merkle et al., 2017), but importantly together with our finding that those cells eliminated in the early mouse embryo have a signature of elevated p53 expression (Lima et al., 2021), suggest that competitive interactions between cells with different levels of p53 expression shape growth during development.

## MATERIALS AND METHODS

### Cell line generation

To generate *p53*^*-/-*^ cells, a p53 sgRNA (5′-GCAGACTTTTCGCCACAGCG-3’) was cloned into lentiCRISPRv2 vector. Viruses were generated by transfecting this vector along with helper plasmids VSV-G and psPAX2 into HEK293T packaging cells. After 48 h, the media from these cells was applied to ESCs with 4μg/ml polybrene. Two rounds of infection, for 4 h and then overnight, were carried out. The cells were then selected using 2μg/ml puromycin and plated at single cell confluency. Clones were screened for loss of p53 protein by western blot.

To generate *p21*^*-/-*^ cells, a p21 sgRNA (5’-GATTGCGATGCGCTCATGGC-3’) was cloned into the px330 vector (Addgene). ESCs were co-transfected with 2μg of this vector and 0,12μg of an hygromycin marker (#631625, TAKARA) using Lipofectamine 2000 (Invitrogen) according to manufacturer’s instructions. The cells were selected using xμg/ml hygromycin and plated at single cell confluency. Clones were screened for loss of p21 protein by western blot.

To generate ^*Bcl2Ind*^ cells, ESCs were co-transfected with 1μg pPB-TRE(3G)-hBCL2-PURO, 1μg pPB-CAG-rtTA(3G)-NEO and 1μg pCMV-PBase using Lipofectamine 2000 (Invitrogen) according to manufacturer’s instructions. The cells were selected using 1μg/ml doxycycline, 2μg/ml Puromycin and xμg/ml Neomycin and plated at single cell confluency. Clones were screened by western blot for BCL2 protein overexpression upon 1μg/ml doxycycline treatment.

### Cell culture

All cells were cultured at 37°C in an atmosphere with 5% CO2. Reagents used for tissue culture were obtained from Invitrogen unless otherwise stated. Mouse embryonic stem cells (ESCs) were cultured on 0.1% gelatin-coated flasks (Nunc, Thermo Fisher) in GMEM containing with 10% (v/v) foetal calf serum (FCS; Seralab), 1X non-essential amino acids, 2 mM L-glutamine, 0.1 mM β-mercaptoethanol and supplemented with homemade leukaemia inhibitory factor (1:1000, LIF). ES cells were routinely dissociated with trypsin and cryopreserved in 10%DMSO in FCS.

### Competition assay

Cells were seeded onto plates coated with fibronectin (Merk) at a concentration of 2.5 × 10^4^ cells/cm^2^ either separately or mixed for co-cultures at a 50:50 ratio (Sancho et al., 2013). Cells were cultured in N2B27 media (Neurobasal media; DMEM F12 media, 0.5 x B27 supplement; 0.5 x N2 supplement; 0.1mM 2-mercaptoetanol; 2mM glutamine; all Thermo Fisher Scientific) during 3-4 days to allow for differentiation. At the indicated time points, the cells were counted using Vi Cell Counter and Viability Analyser (Beckman Coulter) and proportions of each cell type in co-cultures were determined using LSR II Flow Cytometer (BD Bioscience). Sytox blue (Thermo Fisher Scientific) or propidium iodide (Sigma) was used to stain for dead cells.

### Perfusion assay

Cells were seeded into 6-well perfusion plates (AVP011, Reprocell) coated with fibronectin either separately or mixed for co-cultures at a 50:50 ratio in N2B27 media. At day 1, the plate was connected to a bottle with 200 ml N2B27 and perfused using a Watson Marlow 120U Pump with a speed of 9pm. The media circulated from the bottle to the plate, where each well is connected to the next one by a channel, so a unidirectional flow of media is established to each well, and then back to the bottle. The culture was perfused until day 4, when the cells were counted or fixed.

### Fences and conditioned media assays

For Fence assays fences (Aix-Scientifics) were placed in each well of a 24-well plate coated with fibronectin. 0.008×10^6^ cells were seeded in the inner ring and 0.035×10^6^ cells in the outer ring. The fences were removed the following day and the media was replaced every day. At the indicated time points the fences were replaced into the well to count cell numbers using Vi Cell Counter and Viability Analyzer (Beckman Coulter).

For conditioned media assays, 0.08×10^6^ Wild-type or *p53*^*-/-*^ cells were seeded and from day 2 cells were cultured with conditioned media obtained from wild-type or *p53*^*-/-*^ cells cultured separately or from a co-culture. Conditioned media was obtained from the corresponding cell types and was concentrated using Vivaspin 500 centrifugal concentrators (G E Healthcare) according to manufacturer’s instructions.

### Flow cytometry staining

Cells were detached from the plates using accutase (Sigma) and fixed in 7.4% formaldehyde in N2B27 media for 10 minutes. Permeabilization was carried out using ice-cold methanol and cells were blocked using 1% BSA. Cells were then incubated with primary antibody (Cleaved Caspase 3 #9664, Cell signaling,1:200 dilution) for 1 hour at room temperature. After washing, cells were incubated with the secondary antibody (Alexafluor-546/405, Thermo Fisher Scientific, 1:2000 dilution) for 30 minutes at room temperature. Flow cytometry was performed using LSR II Flow Cytometer and analysed using FlowJo software v9 or v10.0.7r2 (BD Bioscience).

### Proliferation assay

Cells were incubated with EDU from the Click-iT kit (Thermo Fisher Scientific) for 2 hours according to manufacturer’s instructions. Cells were then detached from plates using accutase and analysed by flow cytometry using LSR II Flow Cytometer and FlowJo software.

### Stem cells immunofluorescence

For immunostaining ESCs and EpiSCs were fixed for 10min in 4%PFA at room temperature, permeabilised in 0,4% Triton-X100/PBS for 5 minutes at room temperature, blocked in 10%BSA/ 0,1% Triton X-100/PBS and incubated overnight at 4°C in primary antibody diluted in 1%BSA/0,1%Triton X-100 (anti-Cleaved Caspase 3 Asp175, Cell Signalling, 1/100; anti-Nanog (14-5761-80 eBioscience, 1/100, anti ATP-b (Ab14730, Abcam – 1:200). Alexa-Fluor conjugated secondary antibodies (Thermo Fisher Scientific) were used at 1/500 dilution in 1%BSA/0,1%Triton X-100. Cells were mounted for visualization in Vectashield with DAPI (Vector Laboratories). Images were acquired with a Zeiss confocal microscope and analysed with the Fiji software (Schindelin et al., 2012).

### Western blot analysis

Cell lysates were collected in Laemmli buffer (0.05M Tris-HCl pH 6.8, 1% SDS, 10% Glycerol, 0.1% β-mercaptoethanol) and denatured for 10 minutes at 95°C, quantified using BCA quantification (Thermo Fisher Scientific) and resolved using CriterionXT pre-cast gels (BioRad) and transferred to nitrocellulose membranes. Blocking was performed in 5% milk in TBST buffer and primary antibody incubation (p53 #2524, p21, Bcl2, Tubulin) was done overnight at 4°C in TBST containing 5% BSA. Western blot quantification was performed using Fiji software version 2.0.0-rc-49/1.51d. Protein expression levels were normalized to loading control tubulin.

### RNA Extraction and Quantitative RT-PCR

Total RNA was extracted with the RNeasy mini kit (Qiagen) and SuperScript III reverse transcriptase (Thermo Fisher Scientific) was used for cDNA synthesis according to manufacturer’s instructions. Quantitative RT-PCR was performed by amplification with SYBR green Master Mix (Roche). The primers used are listed in Table 1. RNA samples from wild type and mutant clones were collected from 3 independent experiments.

**Table 1:**
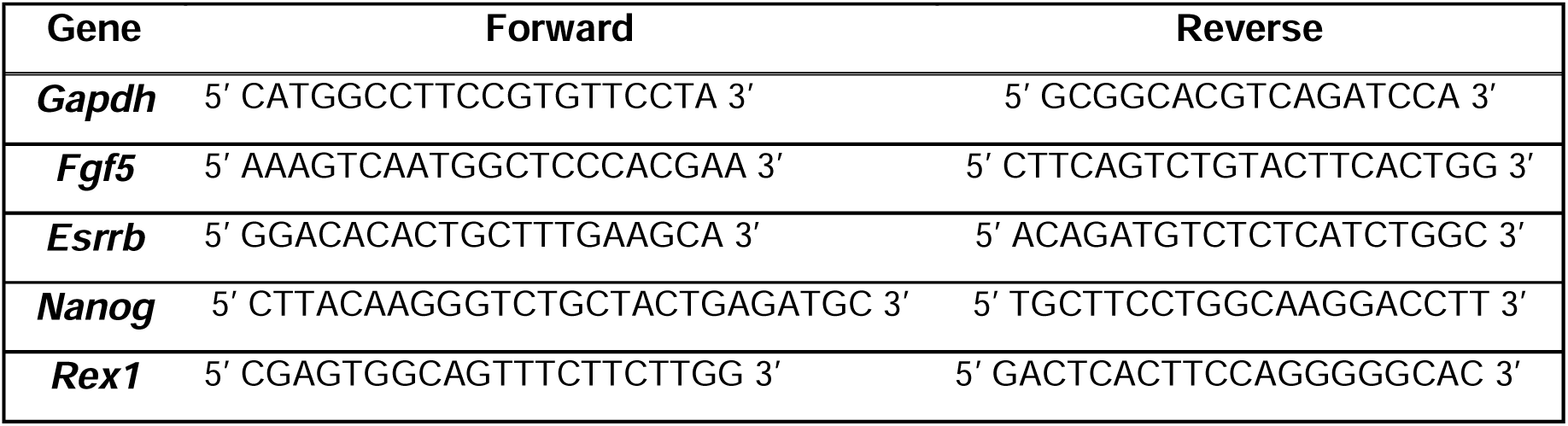
Primers used in the quantitative RT-PCR

### Statistical methods

Statistical analysis was performed using GraphPad Prism v8.0.0 software. Statistical methods used are indicated in the relevant figure legends. No randomization or blinding was used in experiments. Sample sizes were selected based on the observed effects and listed in the figure legends. Statistical significance was considered with a confidence interval of 0.05%; * p<0.05; ** p<0.01; *** p<0.001; **** p<0,0001.

Data obtained from cell competition assays were analyzed by two-way ANOVA, followed by Holm-Sidak’s multiple-comparison test. Data obtained from RT-PCR were analyzed by one-way ANOVA followed by Turkeys post-hoc test. Data obtained from western blot were analyzed with an unpaired two tailed *t*-test.

### Model simulation and Approximate Bayesian inference

Ordinary differential equations where solved with a custom explicit forward Euler method. Parameter distributions compatible with the model were inferred using a Markov chain Monte Carlo implemented in the multiple-try differential evolution adaptive Metropolis algorithm PyDREAM (Shockley et al., 2018). The log-likelihood function for the parameter set *θ* assumed that experimental trajectories deviations (*W*_*exp*_, *P*_*exp*_) followed a Gaussian distribution centred at each timepoint of the observed trajectories (*W*_*model*_, *P*_*model*_) with a width given by the square root of the population size,

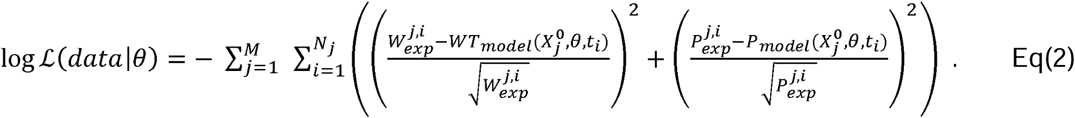

Where the index *j* runs for all the experimental initial conditions 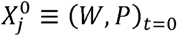; and the Index *I* for all the replicates and their time points *t*_*i*_ Convergence of the posterior parameter distributions was assessed using the Gelman Rubin diagnostic with a threshold *R*_*c*_ =1.2 across 5 different sampling chains.

### Resource competition model

The scenario in which cells compete for an external resource was modelled introducing an additional chemical species *R* that is depleted with cell specific rates *γ*_*W*_ and *γ*_*P*_. The nutrient reduces the intrinisic apoptotic rate of each cell species *δ*_*W*_ and *δ*_*P*_ depending on the characteristic concentrations *r*_*W*_ and. *r*_*P*_.

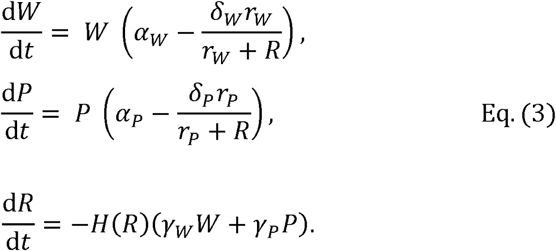

Where *H*(*R*) is the Heaviside step function.

## ACKNOWLEDGEMENTS

We would like to thank all members of the molecular embryology group for their technical and scientific support, especially Dr. Aida di Gregorio and Dr. Ana Lima. We would also like to thank Stephen Rothery for guidance and advice with confocal microscopy. We thank James Elliot and Bhavik Patel from the LMS/NIHR Imperial Biomedical Research Centre Flow Cytometry Facility for support.

## FIGURE LEGENDS

**Figure S1.**
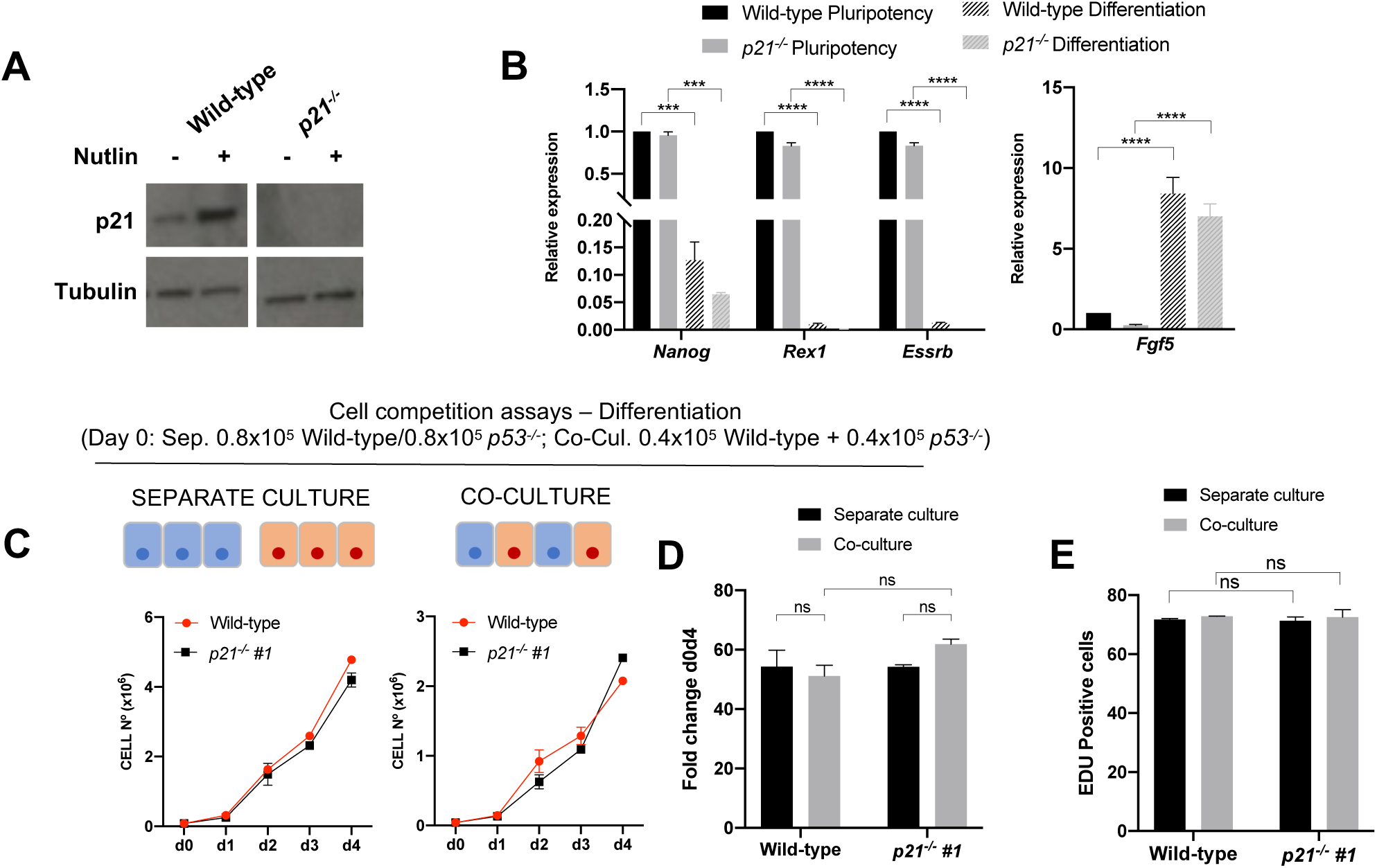
**A**. P21 levels in wild-type and *p21*^*-/-*^ clones untreated or treated with the P53 activator Nutlin-3a for 4h. **B**. Quantitative RT-PCR showing gene expression levels of naïve and primed pluripotency markers in wild-type and *p21*^*-/-*^ ESCs in pluripotency and differentiation culture conditions. Gene expression is normalized against beta-Actin. **C**. Growth curves of wild-type and *p21*^*-/-*^ cells cultured for 4 days in separate or co-culture differentiation conditions. **D**. Fold change in wild-type and *p21*^*-/-*^ cell numbers between day 0 and day 4 when cultured separately or co-cultured. **E**. Percentage of EDU incorporation in wild-type and *p21*^*-/-*^ cells at day 4 cultured separately or co-cultured in differentiation conditions. Data were obtained from three independent experiments and are shown as the mean + SEM (**B**-**E**).

**Figure S2.**
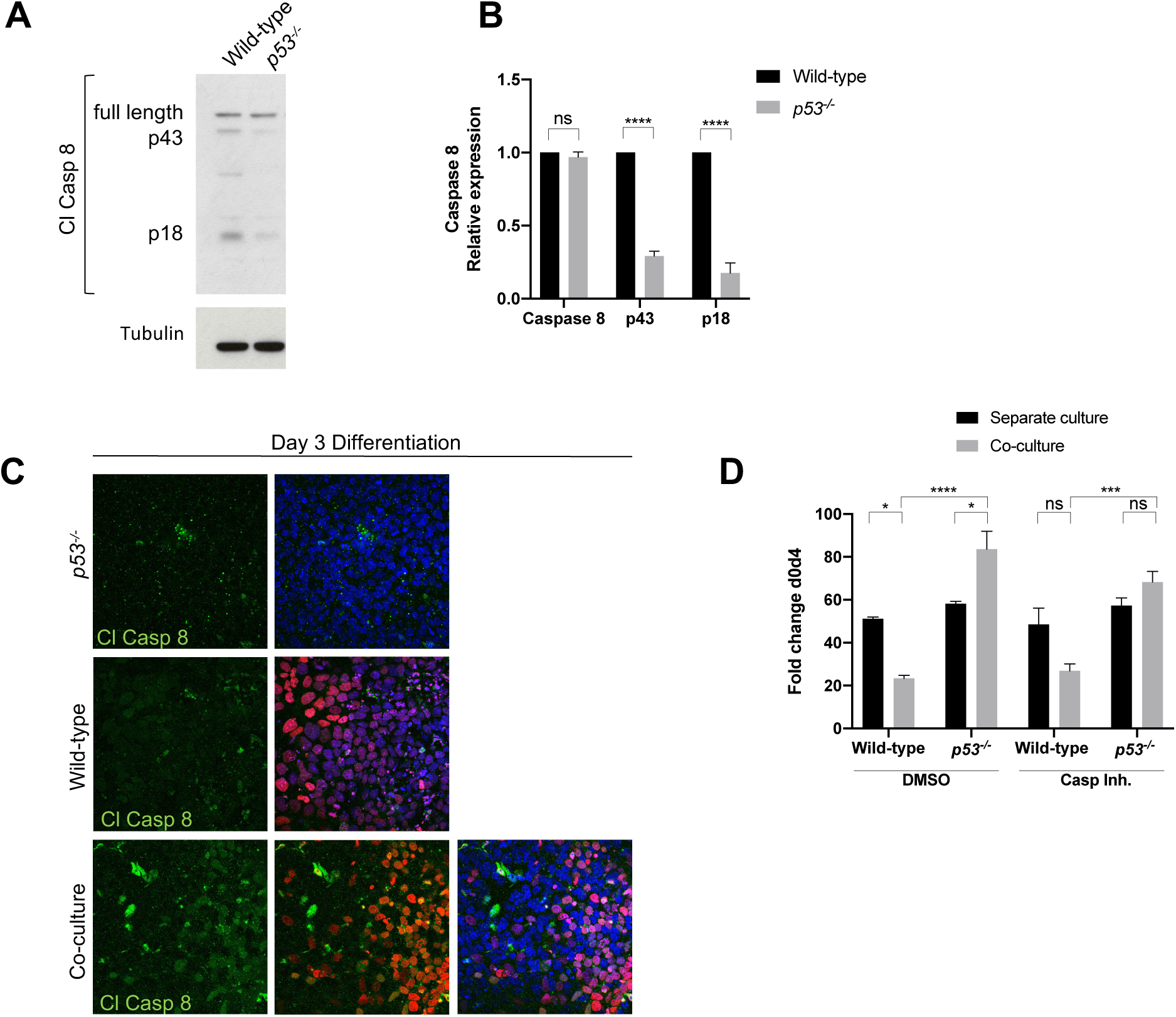
**A**. Levels of un-cleaved Caspase 8 and its cleaved forms p41 and p18 in wild-type and *p53*^*-/-*^ cells cultured separately for 3 days in differentiation conditions. **B**. Quantification of a. **C**. Immunostainings of wild-type and *p53*^*-/-*^ cells cultured in separate conditions or co-cultured for 3 days showing an increase in Cleaved Caspase 8 in wild-type cells. **D** Fold change in wild-type and *p53*^*-/-*^ cell numbers between day 0 and day 4 when cultured in separate conditions or co-cultured in differentiation conditions and treated with DMSO or pan caspase inhibitors from day 3 to day 4. Data were obtained from three independent experiments and are shown as the mean + SEM (**A**-**D**).

**Figure S3.**
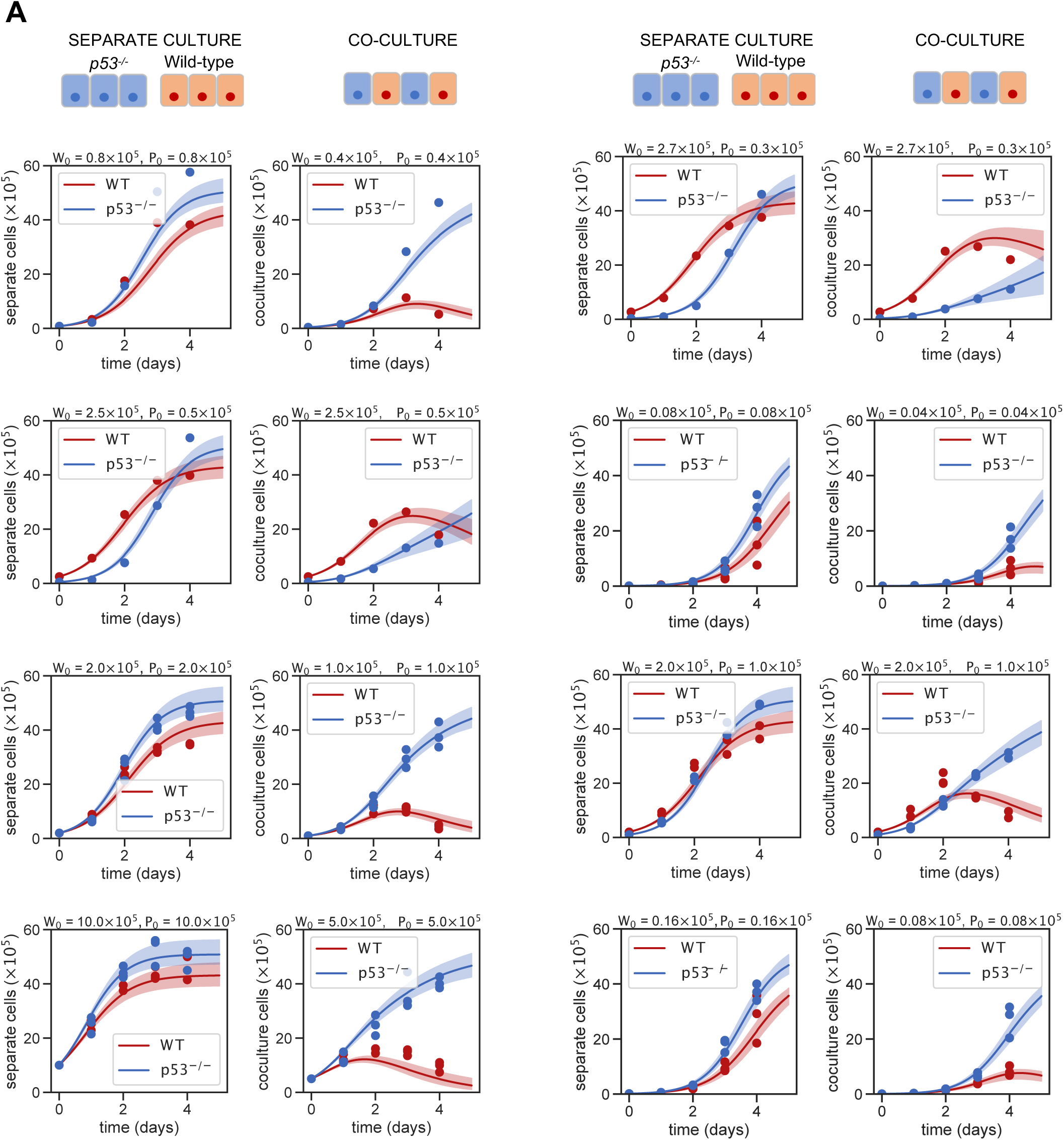
Comparison of the experimental data (one circle per replicate) with the model prediction (lines) for the direct competition model (Eq. 1) for all the different 24 experimental plating conditions used in the inference. Shaded zones show model prediction for the inferred parameter region with likelihood >90% of its maximum. Parameter inference values can be found in Fig. S4.

**Figure S4.**
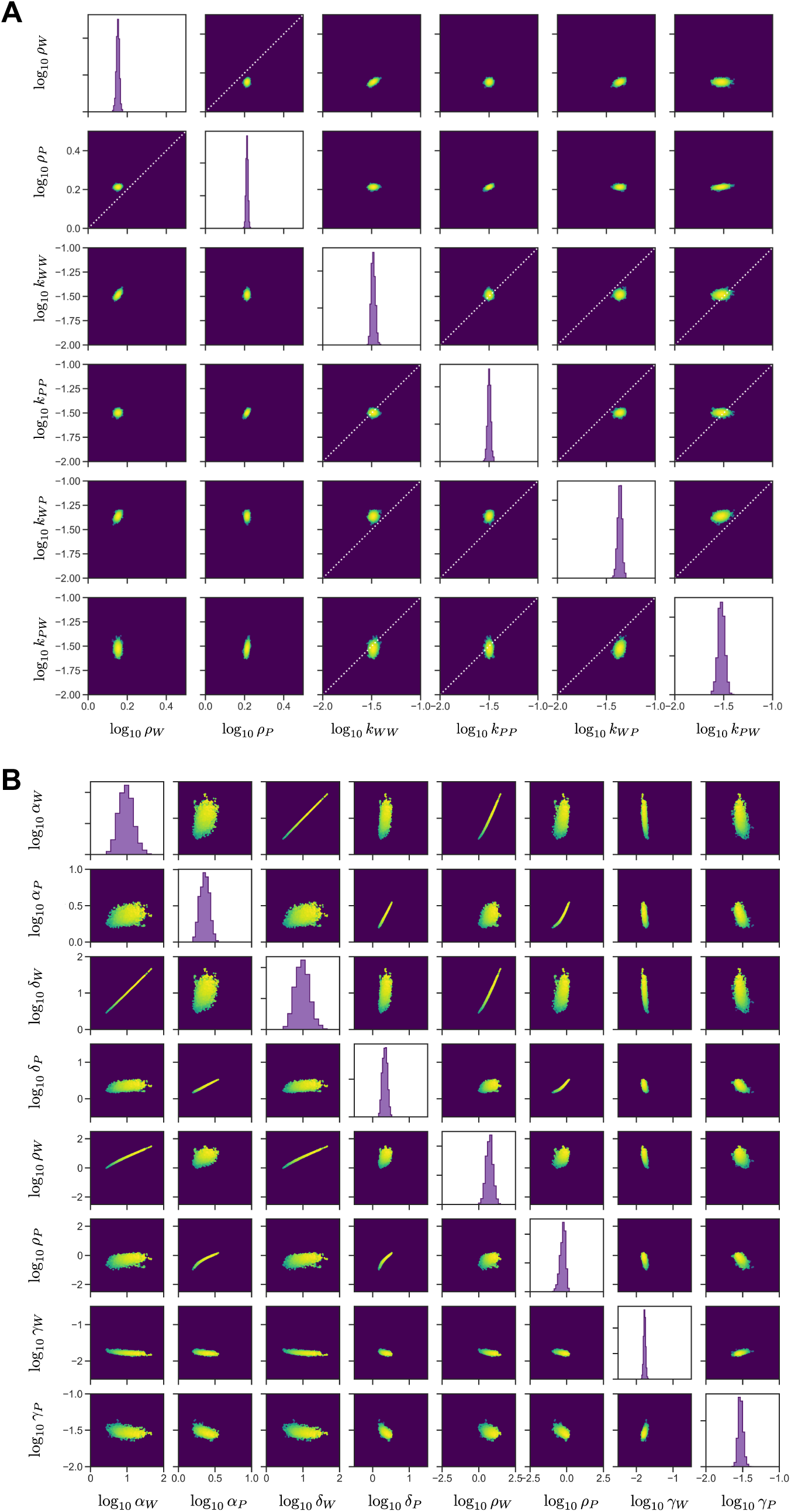
Posterior credibility distributions for the direct competition model (Eq. 1) (**A**), and for the resource competition model (**B**). Diagonal panels show marginal posterior distribution histograms. Off-diagonal panels show the sampled pairwise joint distribution for all the possible parameter pairs. Colour indicates magnitude of the likelihood.

**Figure S5.**
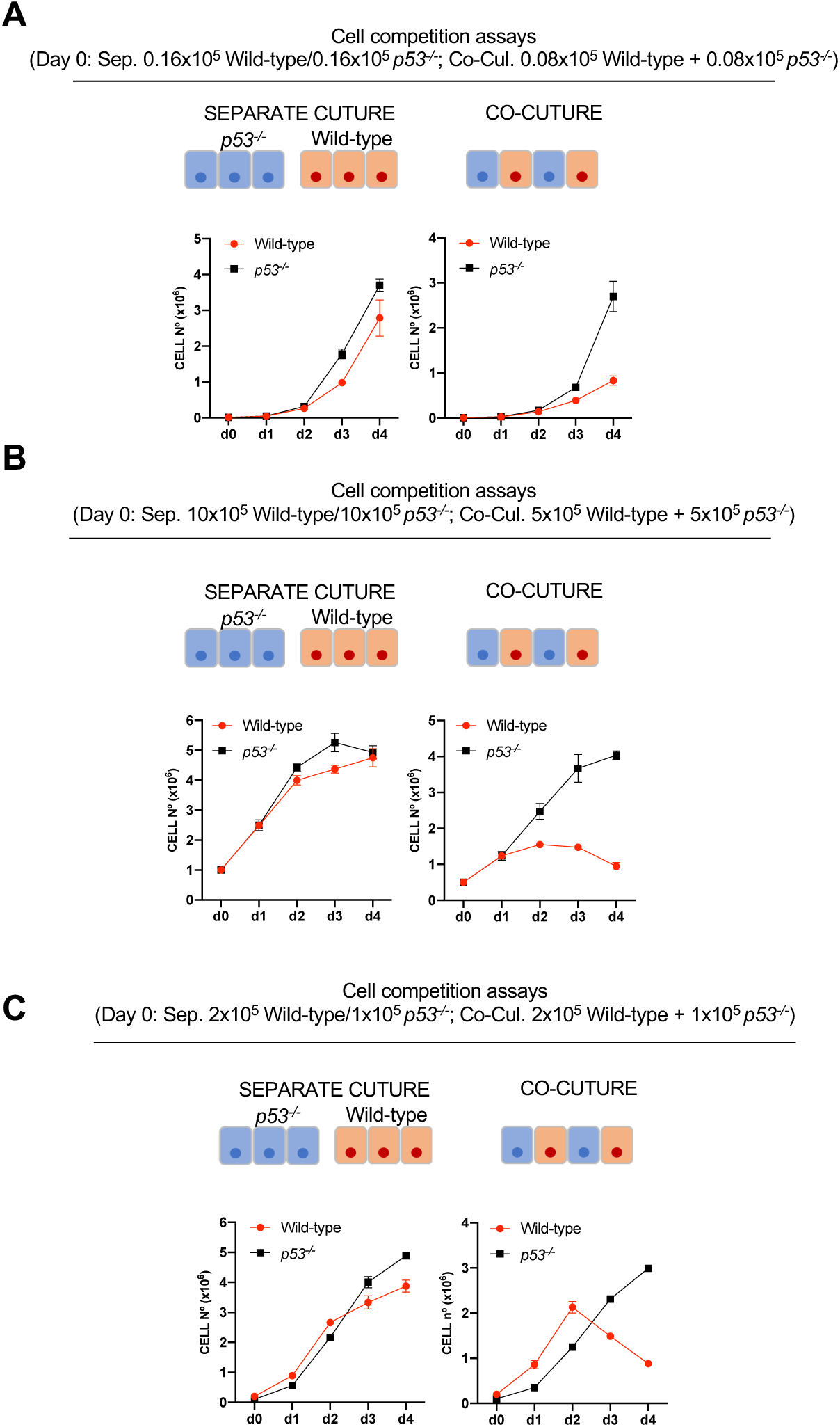
**A**. Growth curves of wild-type and *p53*^*-/-*^ cells cultured for 4 days in separate or co-culture conditions with a starting cell number of 0.016×10^6^ cells seeded per well. **B**. Growth curves of wild-type and *p53*^*-/-*^ cells cultured for 4 days in separate or co-culture conditions with a starting cell number of 1×10^6^ cells seeded per well. **C**. Growth curves of wild-type and *p53*^*-/-*^ cells cultured for 4 days in separate or co-culture conditions with a starting cell number of 0.2×10^5^ wild-type cells and 0.1×10^5^ *p53*^*-/-*^ cells seeded per well. Data were obtained from three independent experiments and are shown as the mean + SEM (**A**-**C**).

